# The promiscuous and highly mobile resistome of a superbug

**DOI:** 10.1101/2021.02.03.429652

**Authors:** Ismael Hernández-González, Valeria Mateo-Estrada, Santiago Castillo-Ramírez

## Abstract

Antimicrobial resistance (AR) is a major global threat to public health. Understanding the population dynamics of AR is critical to restrain and control this issue. However, no study has provided a global picture of the resistome of *Acinetobacter baumannii*, a very important nosocomial pathogen. Here we analyze 1450+ genomes (covering > 40 countries and > 4 decades) to infer the global population dynamics of the resistome of this species. We show that gene flow and horizontal transfer have driven the dissemination of AR genes in *A. baumannii*. We found considerable variation in AR gene content across lineages. Although the individual AR gene histories have been affected by recombination, the AR gene content has been shaped by the phylogeny. Furthermore, many AR genes have been transferred to other well-known pathogens, such as *Pseudomonas aeruginosa* or *Klebsiella pneumoniae*. Finally, despite using this massive data set, we were not able to sample the whole diversity of AR genes, which suggests that this species has an open resistome. Ours results highlight the high mobilization risk of AR genes between important pathogens. On a broader perspective, this study gives a framework for an emerging perspective (resistome-centric) on the genome epidemiology (and surveillance) of bacterial pathogens.

## Introduction

Antimicrobial resistance is a major global menace to humans all over the world. In this regard, the ESKAPE (*Enterococcus faecium, Staphylococcus aureus, Klebsiella pneumoniae, Acinetobacter baumannii, Pseudomonas aeruginosa*, and *Enterobacter* species) group is a significant cause of deaths and burden disease in many countries ^1^. This group of bacteria can easily acquire antimicrobial resistance genes (ARGs). In 2017 the World Health Organization issued a list of bacterial pathogens for which novel drugs are immediately required ^2^. At the top of this list, having the highest priority status (priority 1: critical), was carbapenem-resistant *A. baumannii*. Notably, most *A. baumannii* hospital infections are caused by multidrug resistant (MDR) isolates.

The resistome is the set of ARGs present in a given species or a particular environment ^3^. Over the last decade the resistome of some species and some ecological communities have been studied to an unprecedented detail due to the extensive amount of genetic information provided by genome sequencing. For instance, metagenomics and functional genomics studies have shown that ARGs are frequently found in many different environments ^4-8^. These go from human-made environments, such as hospitals ^5^, to pristine and even isolated environments, such as isolated caves ^7^. Furthermore, many ARGs are globally distributed and are rather diverse. Additionally, population genomics and genomic surveillance studies have been very useful to study the geographical and temporal spreading of important MDR lineages ^9-14^. However, most of these studies have used the core genome as the framework to understand the dynamics of the ARGs. Indeed, ARGs are often just mapped onto the core phylogeny. This strategy can be useful for rather clonal species, such as *Staphylococcus aureus*. Nonetheless, in species with high rates of gene content variation, such as *A. baumannii* ^*15*^, this strategy is bound to be problematic. Furthermore, ARGs are commonly found within mobile genetic elements. Hence, ARGs can be easily moved horizontally within and between species. In this regard, the potential transfer of ARGs between pathogens from different genera is especially concerning in hospital settings ^6^. Therefore, to properly track the transmission dynamics of ARGs, population genomics studies focusing on the ARGs *per se* are required.

Over the last decade, and due to its clinical relevance, a lot of knowledge has been gained for *A. baumannii* through the genome sequencing of many hundred isolates. However, despite the rapid accumulation of genomes, there has not been a single study that has tried to use all this information to comprehensively study the population dynamics of the resistome of this species. Here we produce the most extensive view of the resistome of *A. baumannii*. The data set here amassed has almost 1500 high-quality genomes, covers 42 countries and four decades, and contains 149 lineages (Sequence Types). Our results showed that this is a very dynamic resistome, showing high levels of gene flow within the global population of *A. baumannii*. Furthermore, many ARGs are exchanged with other distantly related bacterial pathogens.

## Results

### The vast accessory resistome of *A. baumannii*

To have the most comprehensive picture of the resistome of *A. baumannii*, we included as many genomes as possible; however, we did pay attention not to include lower quality data. Thus, only genomes showing a high-quality assembly were downloaded (see methods). Furthermore, we included only complete and uncontaminated genomes (completeness ≥ 95% and contamination ≤ 5 %, see methods). We also corroborated that all the genomes belonged to *A. baumannii* via an Average Nucleotide Identity (ANI) analysis; where only isolates having ≥ 95% identity with the type strain ATCC19606 were considered. We kept a total of 1472 genomes for downstream analyses (listed in Supplementary Table 1). This is the most extensive data set ever created for this species: it covered isolates from 42 different countries (Figure 1a), a period of time spanning 76 years (four decades without outliers), and 149 Sequence Types (Figure 1b). Then, we used the Comprehensive Antibiotic Resistance Database ^16^, which is rigorously curated and monthly updated, to catalogue and quantify the ARGs in the 1472 genomes. We found that the average number of ARGs per genome was 29.38; the frequency distribution histogram of ARGs per genome is presented in Figure 2a. The ARGs were classified in 199 ARG families and these covered a wide range of drug classes (see Table 1 and Supplementary Table 2). We noted that no single group was present in all the genomes (see Table 1). Twelve ARG families were present in more than 95% of the genomes. These gene families correspond to the RND efflux pumps, well-known intrinsic resistance genes, and were present in most countries and in almost all the STs (see Table 1). We also found that 26 ARG families (13%) were present in less than 95% of the genomes but in more than 15% of the genomes (see Table 1). Remarkably, almost all of these 38 most frequent ARG families were affected by either recombination or horizontal gene transfer (see Table 1). However, the vast majority of the ARG families (161 groups, 81%) were contained in less than 15% of the genomes (see Supplementary Table 2). These ARG families were present in few countries and few STs (see Supplementary Table 2). Taken together, these results show that a considerable amount of ARGs are found within these genomes. Nonetheless, these ARGs do not belong to the core genome and are not strictly vertically transmitted. Notably most of them are just present in less than 15% of the strains.

**Figure 1.**
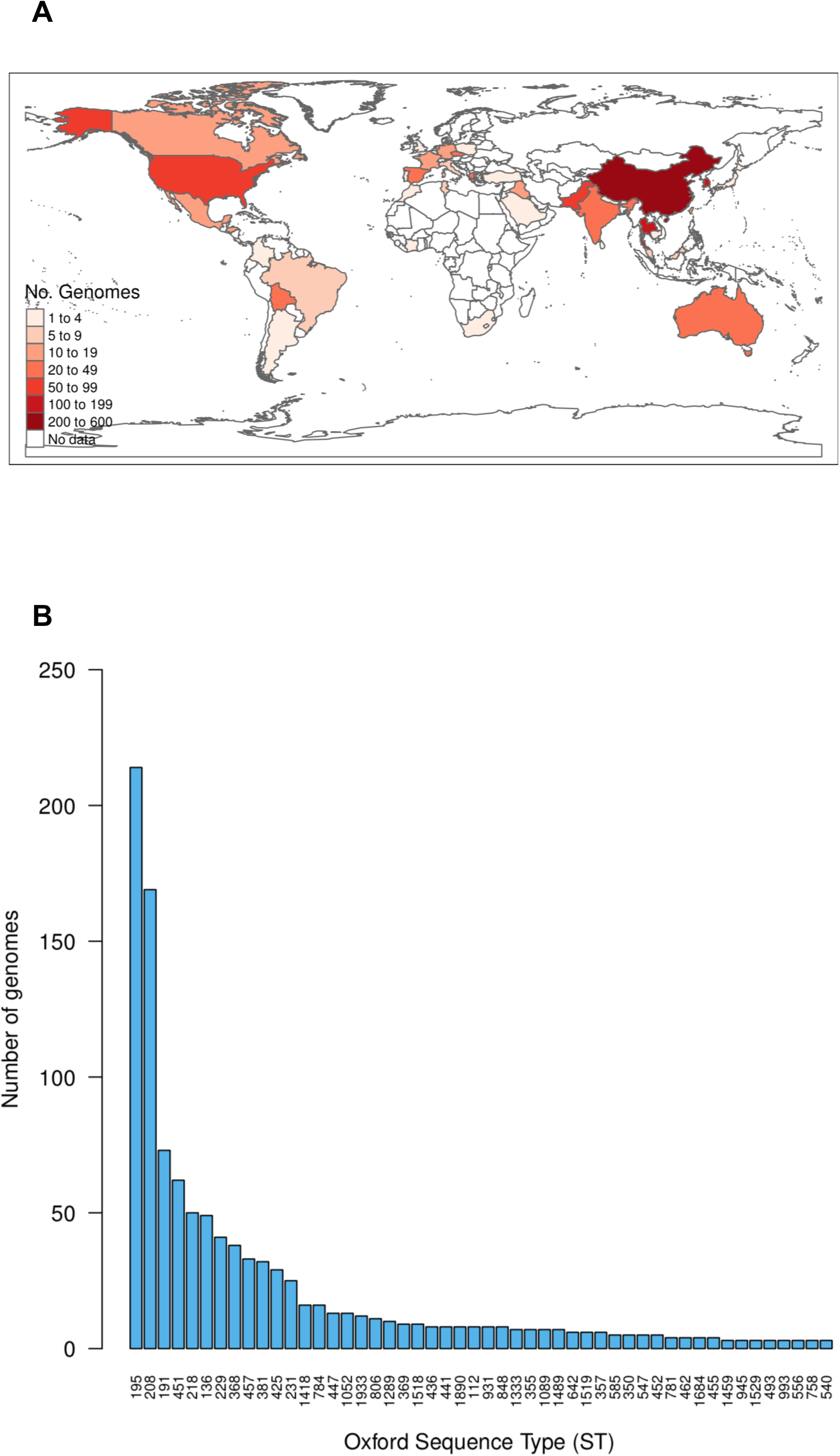
An extensive data set, covering many countries, years, and lineages. (a) Countries from which the genomes were sampled; the color key gives the number of genomes per region. (b) Number of genomes per ST under the Oxford scheme; only STs with 2 or more genomes are shown.

**Figure 2.**
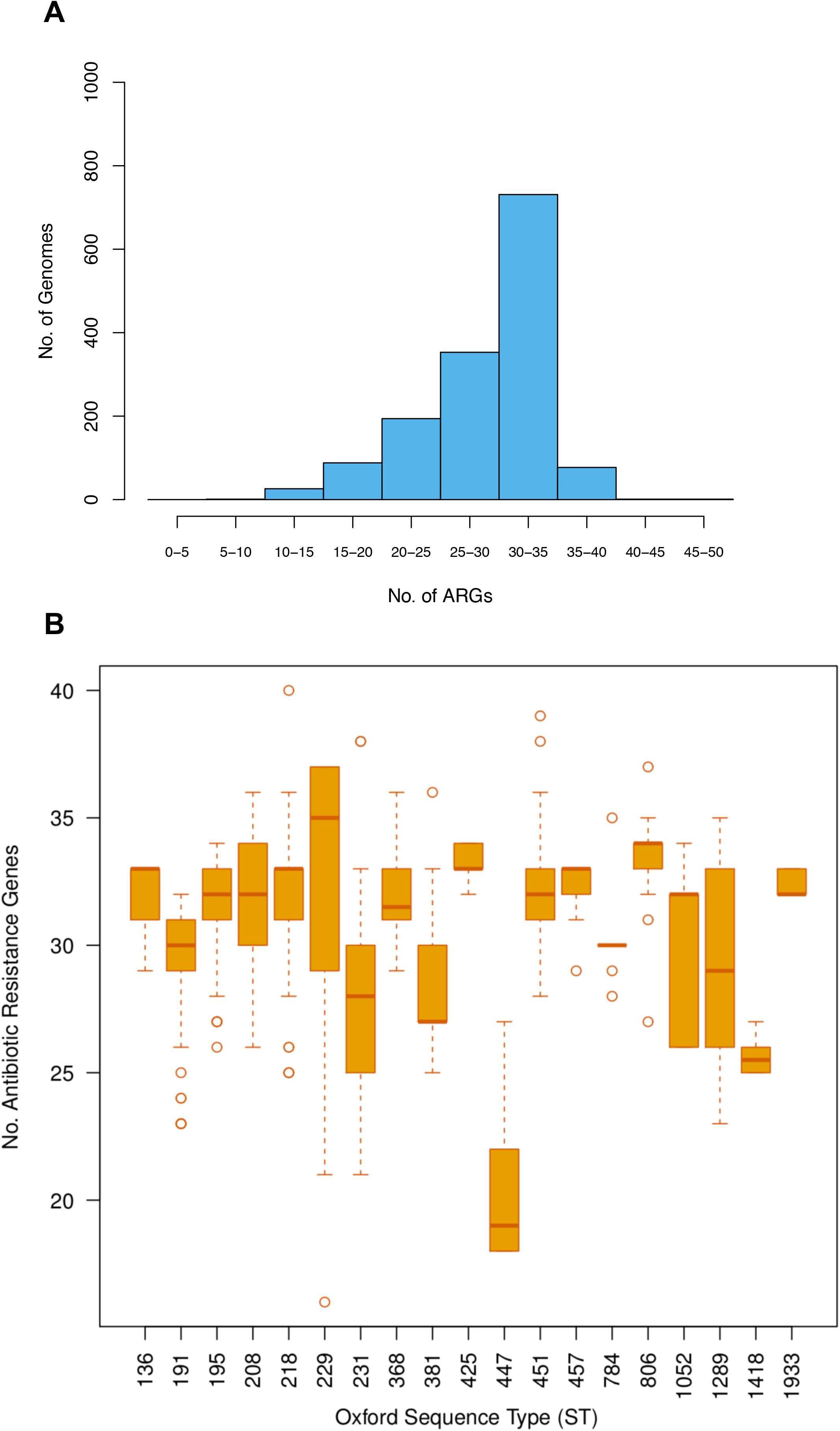
High variation of ARGs. (a) Histogram of the number of ARGs per genome. (b) Boxplots showing the variation in the number of ARGs within and between STs; only STs with at least 10 genomes are shown.

### High gene flow within the species and HGT with other pathogens

The fact that most of the ARGs are just present in some of the genomes implies that ARGs are frequently lost and gained. In accordance with this, we recently showed that gene turnover was of paramount importance in a recently emerged lineage of this species ^15^. To further explore this, we analyze ARG content variation within individual STs; we only considered those STs that had 10 or more genomes per ST. This approach allowed us to analyzed ARGs variation over short timescales. If ARGs were to be only disseminated by clonal expansions (without HGT or gene loss), the amount of ARGs per genome would be the same for a given ST. Contrary to this, Figure 2b clearly shows that there is a huge variation, not only within STs but also between STs (Kruskal-Wallis test, p-value < 2.2e-16). While the ST that had most ARGs was ST229 with an average number of 32 ARGs per genome, the ST with the lowest mean number was ST447 (see Figure 2b) with 20 ARGs per genome. Thus, this high variation in ARG content within STs implies that, even at short timescales, acquisitions and losses of ARGs are very common. Then we went on to look if any of these ARG families have been horizontally transferred very recently (see methods). We noted that 78 ARG families (39%) had identical allelic variants in other bacteria (see Table 1 and Supplementary Table 2).

Notably, most the HGT events were located in other nosocomial pathogens (see Supplementary Table 2 and Figure 3a); *Klebsiella pneumoniae* and *Pseudomonas aeruginosa* are two of the most striking cases. Collectively, these results show that HGT (and gene loss) are of paramount importance not only within the species but also with other (nosocomial) bacterial species.

**Figure 3.**
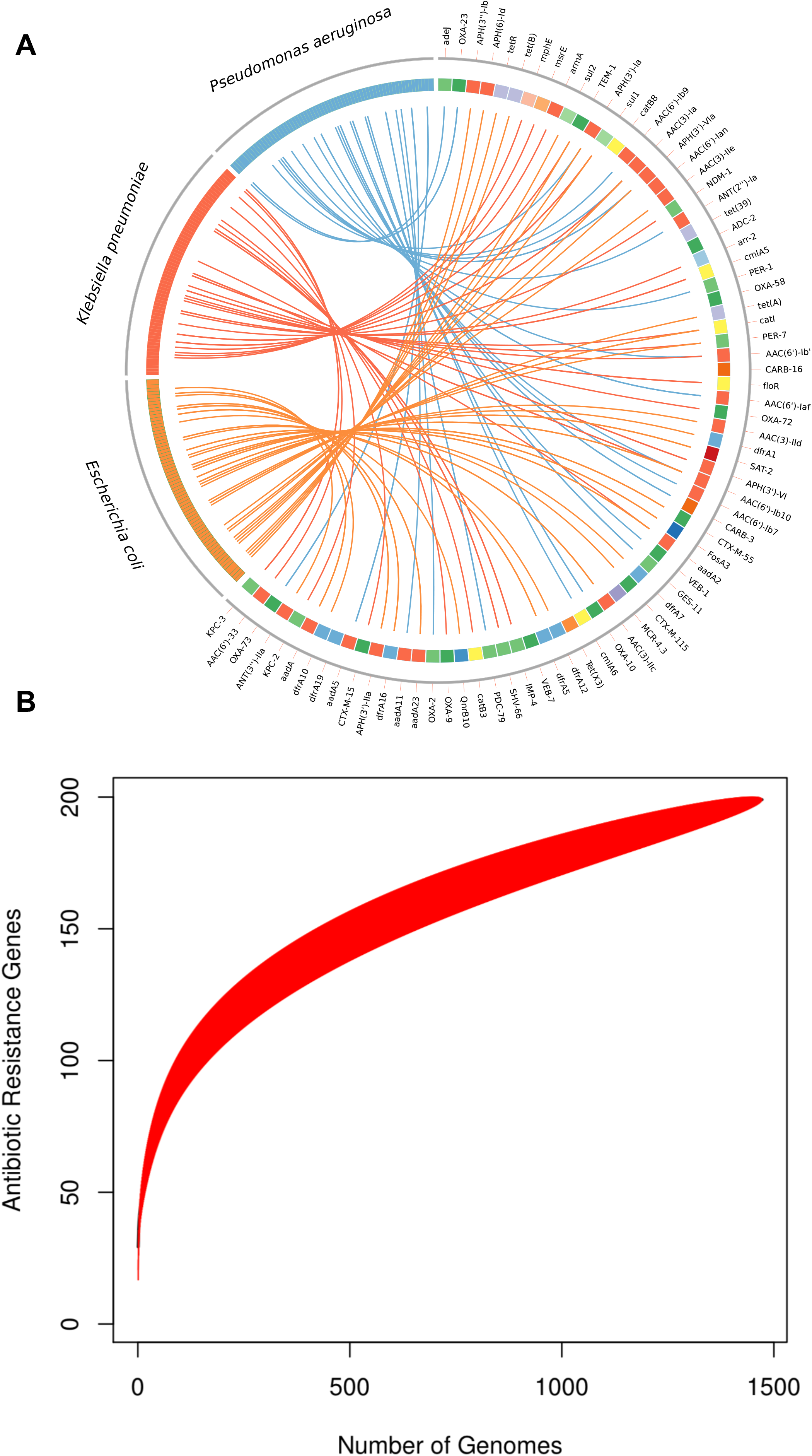
Horizontal transfer of ARGs and an open resistome. (a) Circos plot showing the cases of HGT between ARGs *A. baumannii* (the little squares from 1 to 7 clockwise) and other bacteria (on the left-hand side). Only bacteria (7 to 12 clockwise) involved in 97% of the HGT cases are shown on the left-hand side. (b) Accumulation curve of the number of ARGs as a function of the number of genomes. The red area shows the confidence intervals.

### An open resistome with ARG content structured by the phylogeny

Although ARGs are subject to loss and gain processes, the presence of the ARGs could be structured by the phylogeny. This is because, when considering the accessory genome, HGT between closely related lineages is likely to be more successful; also, the more distantly related two lineages are, the higher the likelihood of gene loss. Thus, to explore this, we conducted pairwise comparisons of all the isolates correlating how similar they were in terms of their ARGs versus their ANI values. We used ANI as a proxy for the phylogenetic relationship of the isolates. To establish how similar the ARGs profiles between the isolates were, we used the Jaccard index. A perfect correlation would imply that the gains and losses of ARGs are shaped by phylogeny. Although not very strong, the Spearman’s correlation was significant (r^2^= 0.57, p-value < 2.2e-16). Thus, there is a correlation between ANI values and similarity in the ARGs profiles, which implied that ARGs content variation correlated with the phylogeny. Finally, we wanted to establish if we were able to sample the diversity of ARGs in this species. Of note, given the very extensive nature of this data set in temporal and geographical terms, we would expect so. We ran an accumulation curve analysis to evaluate this (see methods and Figure 3b). This analysis showed that we have sampled much of the ARG family diversity. However, the curve did not level off and the slope is still very steep in the last section of the curve. Thus, the discovery rate of ARGs is still high after almost 1500 genomes and a lot more ARGs remain to be found. Taken together, these results imply that although gene flow is common within ARGs, ARGs content variation within the isolates is structured to some extent by the phylogeny. Furthermore, this seems to be an open resistome.

## Discussion

Metagenomics studies over the last decade have been of paramount importance to characterize the resistome dynamics in different environments. However, reliable population genomic studies are still very much required to properly describe the transmission dynamics of ARGs both within and between bacterial species. A clear understanding of these transmission dynamics is essential for the effective use of antibiotics. Notably, this will provide useful knowledge to inform public health actions and infection control teams to establish better treatment options. However, there has not been a single study on the resistome of *A. baumannii* at global scale. Here we gathered the most extensive – in terms of geographic and temporal components – dataset ever created for this important nosocomial pathogen. Our analyses underscore a highly mobile and even promiscuous resistome of *A. baumannii*. We found a considerable variation in ARGs profiles among the different lineages (STs). On top of that, we noted that many ARGs have been exchanged with other well-known pathogens.

In some important bacterial pathogens, such as *K. pneumoniae*, MDR isolates (showing many ARGs) are frequently found in high-risk clones. On the contrary, in *A. baumannii* the ARGs are well disperse among the global population and clonal expansion does not seem to be the major force dispersing the ARGs. Previous studies have shown that gene content variation is of paramount importance for this pathogen^15^. Importantly, even very recently emerged ARGs seems to have experienced HGT between different lineages of this species^13^. One of the most relevant findings of our study, from a clinical microbiology point of view, is that *A. baumannii* readily exchanges ARGs with other highly critical pathogens such as *K. pneumonia* or *Pseudomans aeruginosa*. In this respect, a recent study also found many instances of HGT across MDR bacteria from different genera in a single hospital^17^. In connection with the previous points, infection control and detection strategies considering this pathogen should be focused on the ARGs, rather than on particular lineages, to prevent the transmission of such genes. Thus, the risk of ARGs across important nosocomial pathogens should be considered of paramount importance not only from a microbial genomics point of view but also from infection control perspective. Finally, and connected with the previous point, we observed that, notwithstanding the vast data set we used, our accumulation curve analysis implies that we did not sample the full diversity of ARGs within *A. baumannii*. Thus, very likely this species has an open resistome; given the tendency of this species acquires ARGs from other bacteria an open resistome is not actually unexpected.

One of the limitations of our study is that we underestimated the rate of HGT with other bacteria. This is because we only considered very recent HGT events, *i*.*e*., identical gene sequences in different bacteria. Clearly, the rate of HGT is bound to be considerably higher, if HGT events of different ages are to be included in the estimation. Another limitation of our study is that our analyses were based on a genetic definition of antibiotic resistance^18^. We acknowledged that further studies considering also microbiological and clinical definitions of antibiotic resistance are required to fully grasp the complex issue of antimicrobial drug resistance. Nonetheless, understanding the populations dynamics of the ARGs is crucial for the tracking and surveillance of ARGs not only in the clinic but also in non-clinical settings. In this regard, we recently show that environmental isolates of *A. baumannii* can be an important source of ARGs^19^. Without doubt, the transmission dynamics of ARGs can be used to implement *ad hoc* infection-control actions within different health systems all over the world. Thus, similar studies are required for many other members of the ESKAPE group, or any other relevant human pathogen for that matter, to properly tackle the major problem of antimicrobial drug resistance.

## Methods

### Genomes and quality check

Genomes sequences were downloaded in early March 2020 from the Refseq NCBI Reference Sequence Database. Only genomes with reliable quality genome assembly status were considered; thus, only genomes showing an assembly level of “complete genome”, chromosome and scaffold were included in the final data set. We also used CheckM^20^ to evaluate the completeness and contamination of the genomes and only genomes showing 95% (or higher) of completeness and equal or less than 5% of contamination were considered for downstream analyses. For consistency all the genomes were re-annotated with PROKKA^21^ and the final list of the genomes included is provided in Supplementary Table 1.

### Average nucleotide identity analysis and ST assignation

To make sure that all genomes belong to *Acinetobacter baumannii* and to evaluate how similar were among them, we conducted an Average Nucleotide Identity (ANI) analysis using OrthoANI^22^. Using pairwise comparison, every genome was compared to rest of the genomes and to the type strain ATCC 19606 (Biosample accession number SAMN13045090). We kept only the isolates that had an ANI value of 95% or higher versus the type strain. We downloaded the allelic variants for profiles of all the ST described thus far for both the Oxford and the Pasteur schemes from the PubMLST database^23^. Then, we used blastn (requiring 100% identity) to assign an allelic profile to each genome considering both schemes. Given that the Oxford MLST scheme has some issues with paralogy in the locus *gdhB*, we processed the data as in ^23^ to discard the paralogous genes causing issues.

### Antibiotic resistance gene prediction and pangenome analysis

We employed the CARD database^16^ the infer the ARGs in all the *A. baumannii* genomes. We used the Resistance Gene Identifier tool from CARD and only “Perfect” and “Strict” cases were considered but we also required a ≥ 90 % coverage between the query and the target. Antibiotic drug class were also determined through CARD. Then, the ARGs were assigned to the different types of genome categories as follows: core genome, genes in 100% of the isolates; soft core genome, genes between 95% and less than 100% of the isolates; shell genome, genes in less than 95% but greater than 15 % of the isolates; and cloud genomes, genes in less than 15% of the isolates. The ARGs were grouped into ARG families using the CARD classification.

### Recombination analysis, horizontal gene transfer with other species, and accumulation curve analysis

Recombination analysis were run on all the ARG family groups that had at least 3 sequences and which shared at least 80 % of coverage among all the sequences. We used the PHIpack program^25^ that implements the “Pairwise Homoplasy Index” test (PHI test) to detect recombination signals within the ARG families. To infer recent events of HGT between the ARGs in *A. baumannii* and other bacteria, we used blastn (with an e-value of 1e-50) to search these ARGs against the “nt” database from the NCBI. Off note, as we wanted only very recent HGT events, we considered only those hits that were 100% identical (both in the coverage and in the % of identity) to the query. As query could have identical hits from several species, we chose that species that had highest number of identical hits. We used the Vegan R library^26^ to run a rarefaction curve of the number ARGs as a function of the number of genomes. We employed the “specaccum” function for this, setting the method “rarefraction”. The visualization tool Circos ^27^ was employed to illustrate the HGT cases between the ARGs in *A. baumannii* and the other three major bacteria contributors.

## Acknowledgments

We are very grateful to Alfredo José Hernández Álvarez and Victor Manuel del Moral Chávez for installing several of the programs employed in this study. SCR is grateful to his friend Timothy Read for reviewing the manuscript.

## Authors’ contribution

SCR conceived and supervised the study. VME conducted preliminary analyses of the antibiotic resistance gene prediction. ILHG carried out most of the analyses. SCR wrote the manuscript and all the authors approved it.

## Conflicts of interest

All the authors report no potential conflict of interest.

## Funding

This study was supported by supported by “Programa de Apoyo a Proyectos de Investigación e Innovación Tecnológica” (grant no. IN206019) and “Consejo Nacional de Ciencia y Tecnología” (CONACyT) Ciencia Básica 2016 (grant number 284276) to SCR. VME is a PhD student from the Programa de Doctorado en Ciencias Biomédicas, Universidad Nacional Autónoma de México, and is supported by a CONACyT postgraduate fellowship. ILHG acknowledges support from UNAM Dirección General de Asuntos de Personal Académico postdoctoral fellowship funding.

## FIGURES AND TABLE

**Table 1 – The most frequent ARGs found in the species**

Information for the most frequent ARGs is provided. These ARGs were present in more than 100 genomes. The number of genomes, countries and ST in which every ARG was present is reported. In bold are those ARGs that had recombination signals and the superscript ^**H**^ means that identical allelic variants were found in other bacteria.

## SUPPLEMENTARY MATERIAL

**Supplementary Table 1**

List of the genomes along with their metadata employed in this study.

**Supplementary Table 2**

Information of all the 199 ARGs found in this study. The number of genomes, countries and STs are provided; as well as whether the ARG was subject to HGT.

## Notes

### Competing Interest Statement

The authors have declared no competing interest.

